# Inferential Learning of Serial Order of Perceptual Categories by Rhesus Monkeys (*Macaca mulatta*)

**DOI:** 10.1101/102897

**Authors:** Natalie Tanner, Greg Jensen, Vincent P. Ferrera, Herbert S. Terrace

**Author notes:** Corresponding Authors: GJ, HST, **Submitting Author:** Herbert Terrace, Columbia University, 1190 Amsterdam Ave. MC 5501, 406 Schermerhorn Hall New York, NY 10027.

## Abstract

Category learning in animals is typically trained explicitly, in most instances by varying the exemplars of a single category in a matching-to-sample task. Here, we show that rhesus macaques can learn categories by a transitive inference paradigm in which novel exemplars of five categories were presented throughout each training session. Instead of requiring decisions about a constant set of repetitively presented stimuli, we studied the macaque’s ability to determine the relative order of multiple exemplars of particular stimuli that were rarely repeated. Ordinal decisions generalized both to novel stimuli and, as a consequence, to novel pairings. Thus, we showed that rhesus monkeys could learn to categorize on the basis of implied ordinal position, and that they could then make inferences about category order. Our results challenge the plausibility of association models of category learning and broaden the scope of the transitive inference paradigm.

**Significance Statement:** The cognitive abilities of non-human animals are of enduring interest to scientists and the general public because they blur the dividing line between human and non-human intelligence. Categorization and sequence learning are highly abstract cognitive abilities each in their own right. This study is the first to provide evidence that visual categories can be ordered serially by macaque monkeys using a behavioral paradigm that provides no explicit feedback about category or serial order. These results strongly challenge accounts of learning based on stimulus-outcome associations.

---

Since the discovery that pigeons could be trained to peck at photographs that only contain people (Herrnstein & Loveland 1964), an extensive literature has demonstrated an animals’ ability to categorize a wide variety of stimuli, e.g., faces (Marsh and MacDonald 2008), plants, and animals (Roberts 1996), man-made objects (Bhatt et al. 1988), and even paintings (Watanabe 2013). Freedman and Miller (2001) found that primates could categorize computer-generated, systematically morphed images of cats and dogs. Activation in the lateral prefrontal cortex correlated with the stimulus category even when stimuli were assigned to new categories (Freedman and Miller 2001).

Although many studies have demonstrated that animals can categorize stimuli, relatively little work has been done showing how categories are used in other cognitive tasks. Can animals, for example, treat categories as though they were informative stimuli, signaling appropriate behavior in a cognitive task?

## Categorical Serial Learning

Altschul and colleagues (2016) demonstrated that rhesus macaques can not only distinguish between four simultaneously-presented categories of stimuli, but that they can also learn their serial order using a variant of the Simultaneous Chain task (Terrace 1984, 2005). This suggests that animals can not only learn to identify categories but they can also manipulate categories in the same way they can manipulate single stimuli. That is, they applied judgments of list position to entire classes of stimuli.

Transitive inference (TI) is another paradigm for studying serial learning: the ability to learn the relative order of a set of items. TI has been demonstrated in many species, including monkeys (McGonigle & Chalmers 1992), pigeons (Lazareva & Wasserman 2006), crows (Lazareva et al. 2004), and even fish (Grosenick et al. 2007) (for review, see Vasconcelos, 2008; Jensen, 2017). At its most abstract, TI involves maintaining a representation of the relative order of list items. Following training using only adjacent pairs, above-chance performance on non-adjacent pairs demonstrates that subjects were capable of TI (McGonigle and Chalmers 1977, Jensen et al. 2013). While an animal’s ability to learn a TI task and to categorize are well established, their ability to do both simultaneously has yet to be shown. Here, we assess the ability of rhesus macaques to learn category membership of stimuli that change on every trial, even as they learn the order of those categories and ultimately perform TI for critical test pairs.

Our Category TI task follows the format of a traditional TI procedure (train on adjacent pairs, test on all pairs), but does so with stimuli that change for *every* trial. Subjects used trial and error to learn the category order of stimuli belonging to five categories by trial and error: birds, cats, flowers, people, and hooved mammals. Each trial begins with the presentation of two randomly selected pictures, drawn from a pool of 1000 images for each of the five image categories. Because the images included a range of related species photographed under varying conditions, subjects had to rely on category membership rather than their memory of specific stimuli.

Subjects learned all of the categories while also learning the list order. They had no prior exposure to categorization tasks generally, or to any of the exemplar stimuli used for those categories. After subjects were tested for TI with one stimulus order, the same categories were trained again, this time using a different category order. During the course of the experiment, subjects had to learn to sort the five categories into four different orderings. Given the size of the stimulus sets and the lack of prior training, a demonstration of TI under these conditions would show that perceptual categories can be deployed and represented in the same flexible fashion as the constant stimuli that are normally used in TI tasks. It would also show that subjects can learn to categorize images without an initial training procedure designed solely to train category membership (e.g. match-to-sample).

## Methods

### Subjects

Subjects were two adult rhesus macaques (*Macaca mulatta*), N and O. Both had prior experience with transitive inference procedures, but neither had any experience with categorizing pictorial stimuli. The early phases of the experiment were the subjects’ first exposure to these five categories.

The subjects were individually housed in a colony room containing approximately two-dozen macaques and performed the experimental tasks in their home cage. The subjects were trained five days a week, completing one session each day. To increase their motivation to perform the task for fluid reward, the monkeys were put on fluid restriction (300mL per day) two days prior to the first day of testing. Depending on task performance, subjects earned between 200mL and 300mL a day performing the task. Most days, subjects earned their entire fluid ration performing the task. This was supplemented as needed after the experimental session ended to meet the minimum. Each monkey received a set amount of biscuits each morning prior to testing. Fruit was distributed following testing.

The study was carried out in accordance with the guidelines provided by the *Guide for the Care and Use of Laboratory Animals* of the National Institute of Health (NIH). This work, carried out at the Nonhuman Primate Facility of the New York State Psychiatric Institute, was overseen by NYSPI’s Department of Comparative Medicine (DCM) and was approved by the Institutional Animal Care and Use Committees (IACUC) at Columbia University and NYSPI.

### Apparatus

The apparatus used for this study was an in-cage testing device with a touch-screen tablet and a fluid delivery system comprising a 1L calibrated reservoir and a solenoid valve. The solenoid valve was controlled by the tablet computer via an Arduino Nano interface. Each correct response delivered 0.25mL of water through a spigot below the touch-screen. The entire testing device fit snugly and securely into the doors of the monkey’s home cages. The tablet had a 10.1” HD display, operated at 1266 × 768 pixel resolution, and used capacitative multitouch inputs. All tasks were programmed in JavaScript and run in a Google Chrome browser window under a Windows 8.1 operating system.

All stimuli used in the experimental tasks were 250 × 250 pixel images presented randomly on the left and right-hand sides of the tablet’s screen. Between trials, a solid blue square was presented at the center of the screen. Touching it initiated a new trial. This focused the subject’s attention and directed the subject’s hand towards the center of the screen to reduce response bias.

### Stimuli

Stimuli were selected from five categories: Birds, Cats (including both housecats and large predatory cats), Flowers, People, and Hoofed Mammals (the last being a mix of sheep, cows, horses, and goats). Other than people, each category comprised a variety of species photographed under varying conditions. For each category, subjects were exposed to 1,000 different stimuli. It was therefore highly unlikely that subjects would see the same image more than once within an interval of several hundred trials. Stimuli from the first four categories were previously used by Altschul et al. (2016) with a different set of subjects.

### The Transitive Inference Procedure

During training, subjects were provided with incomplete information about list order. It was, however, possible for them to infer the relative ordinal position of each item. Consider, for example, a list of arbitrarily selected stimuli (A, B, C, D, & E) in which the order was determined by the experimenter and unknown to subjects. On each trial, subjects were presented with pairs of items. A response to the item from the earlier list position was always rewarded. If, for instance, the order was ABCDE and the pair BC was presented, a response to B was rewarded because it came first. If, however, the pair AB was presented, the subject had to choose A to receive a reward. Following training on adjacent items (AB, BC, etc.), the critical question is whether subjects can infer the correct choice when presented with non–adjacent items that they had never previously seen (e.g. AC).

Each session, subjects completed up to 1,000 trials of a transitive inference (TI) task (cf. Figure 1) by touching stimuli on the tablet to earn water rewards. Each of two images presented during a trial had an associated “list rank” that was not explicitly communicated to the subjects. The image with the lower rank (i.e. earlier in the list) was always correct, and selecting correct items yielded a reward of 0.5mL of water. Image ranks ranged from 1 to 5. Thus, subjects were effectively asked to discover the order of a five-item list (denoted as *ABCDE*) by pairwise trial and error (see Jensen et al. 2013, 2015) in a procedure in which the exemplars of each category were selected at random and seldom repeated.

**Figure 1:**
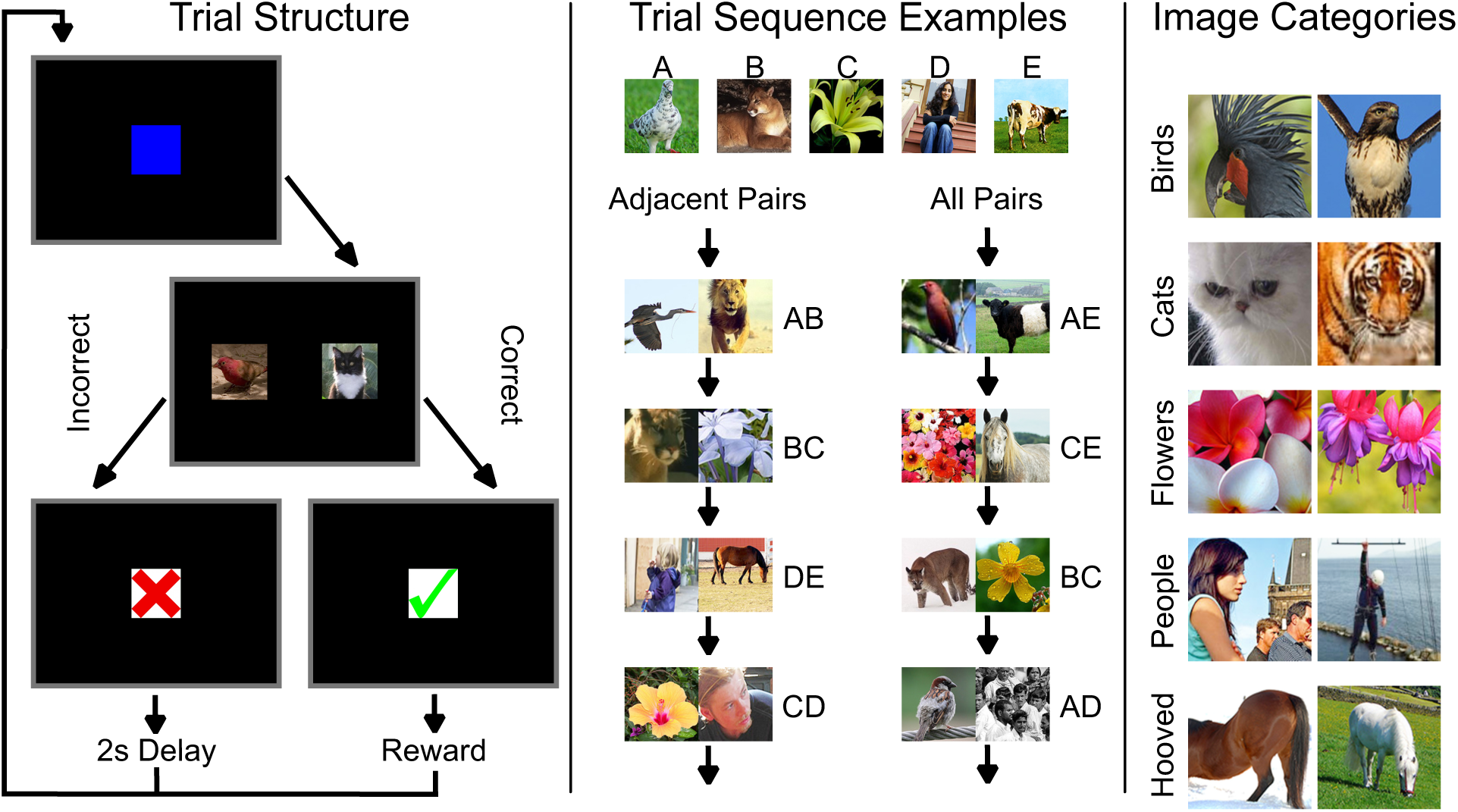
Procedure for the categorical TI task. **Left.** Trial structure for any single trial of the task. Subjects must touch a blue square to begin the trial, which is immediately replaced by two images. If a correct response is made, subjects see a green check mark and are immediately given a fluid reward. If an incorrect response is made, subjects see a red X, followed by a black screen for 2 seconds. Following feedback, the next trial begins with the start stimulus. **Middle.** Each phase of the experiment made use of a consistent category sequence (in this case, birds-cats-flowers-people-hooved). The stimuli themselves, however, were drawn at random from the image back during every trial. During adjacent-pair trial (using only AB, BC, CD, and DE), the identity of the stimulus changed for every trial, even when the same category appeared in two consecutive trials. The left-right position of stimuli was also counterbalanced. This was also the case during all-pairs sessions, which intermixed all possible stimulus pairings. **Right.** Two exemplars each from the five stimulus categories used in the experiment. In all categories, an effort was made to include category members from multiple distances and angles, with a mixture of both solitary and group photos, as well as both color and black-and-white. This stimulus diversity was intended to reduce subjects’ reliance on specific discrete features as categories cues.

Unlike traditional TI tasks, a particular rank was not associated with a single static image. Instead, as described above, rank was associated with a stimulus category. Every time a subject saw the pair *AB*, it consisted of a different random pair of images from categories *A* and *B* than those shown in the previous pairing of A and B. This meant that subjects could not solve the task by learning the order of specific stimuli. Because the images included a range of related species photographed under varying conditions, subjects had to generalize their understanding of one image of a bird and one image of a cat and understand, for example, that all birds come before all cats. Since subjects had no experience with these categories, they had to learn them at the same time they were learning list order. In this respect, our procedure deviated from the typical matching-to-sample or match-to-category procedures used to study concept-formation (Freedman and Miller, 2001; Bodily et al. 2008).

To test subjects’ knowledge of TI, their initial training was limited to adjacent pairs (*AB, BC, CD, DE*). During such training, A is always rewarded, E is never rewarded, and all other stimuli are rewarded half the time. B, for example, is correct when paired with C, but incorrect when paired with A. Its expected value is therefore 0.5. Once subjects performed above chance on such pairings, they were tested on a “critical test pair”, e.g., BD. Because B and D each have an expected value of 0.5, associative models predict performance no better than chance. Contrary to this prediction, subjects across many species routinely favor B, thereby displaying TI, despite B and D having similar reward histories. After at least six sessions of adjacent pair training, subjects were exposed to all ten possible stimulus pairings. Knowledge of TI would be demonstrated if subjects performed at a greater than chance level on the critical pair BD.

The *symbolic distance effect* is a robust feature of TI performance. Stimulus pairs that are more widely separated in the list show higher levels of accuracy than those that are closer together (D’Amato & Colombo 1990, Treichler et al. 2007). Given our “train-adjacent-test-nonadjacent” task design, a symbolic distance effect observed at the *start* of each all-pairs block of sessions would be difficult to explain using associative models, as would above-chance performance on critical test pairs. Such effects are instead consistent with a strategy that relies on the comparison of relative ordinal or spatial position, as widely-spaced items should be easier to discriminate than closely-spaced items.

After an initial transfer from adjacent pairs to all pairs with respect to a particular list order, subjects were again presented with adjacent pairs, this time using a different ordering. They repeated the adjacent-pair-training, all-pair-testing design for three more phases, yielding a total of four different category orders. The order for Phase 1, representing the adjacent and then all-pairs testing, was ABCDE with BD being the key pair for evidence of TI. The order for Phase 2 was DBCEA with BE being the key pair. The order for Phase 3 was AECBD with EB being the key pair. The order for Phase 4 was EDCBA with DB being the key pair. Due to scheduling conflicts and technical difficulties subjects were not run for the same number of sessions: N completed 80 sessions total, whereas O completed 60 sessions. Both subjects consistently completed three sessions before and after each transition.

## Results

We achieved both of our goals by showing that rhesus macaques could, in the context of the TI paradigm, learn (1) to simultaneously categorize photographs from five categories without prior matching-to-sample training and (2) the ordinal position of those categories in an implicitly defined list. Behavior was modeled using logistic regression, building on the method described Jensen and colleagues (2013). The probability of selecting the correct stimulus on trial *t* during a particular session is given by *p(t)*, which was fit according to the following function:

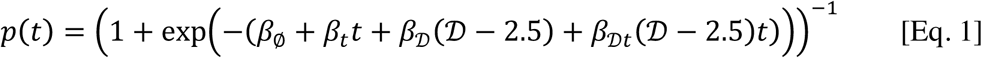

Here, *t* refers to the trial number, beginning with zero; consequently, *β_ø_* is the intercept term, and *β_t_* is the slope as a function of time. *D* refers to the symbolic distance between the list positions of the stimuli (for example, for an adjacent pair, *D*=1). Since the maximum value of *D* is four (in the case of pair *AE*), subtracting 2.5 from *D* in the analysis centers it. As a result, *β_t_* provides an estimate of improvement in performance overall, and *β_D_* represents the *differential* performance that results from the symbolic distance effect. *β_Dt_* represents the interaction between overall learning and the symbolic distance effect. A more compressed version of Equation 1 is to simply report it as the logistic function:

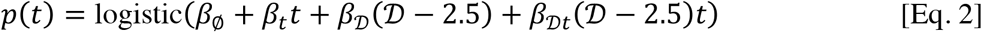

A different logistic regression was performed for each subject during each session because subjects were presented with the same stimuli over multiple consecutive sessions. This allows a distinction to be made between behavior during learning and behavior when performance reached ceiling (which, in macaques, is consistently below perfect accuracy). Models were fit using the Stan language (Carpenter et al., In Press). To facilitate continuity from one session to the next, model estimates for a subject’s performance at the *end* of each session acted as a regularizing prior on that subject’s performance at the *beginning* of the following session. In transitions between phases, earlier performance was not used as a prior. For details, see the electronic supplement.

Figure 2 presents the estimated mean probability of a correct response (combining both subjects) during the first and last three sessions of each phase. Adjacent pairs are plotted in black, while the 80% credible interval for those estimates is shown by the shaded regions. Consistent with the past literature (Terrace 2003, Lazareva 2006, D’Amato 1990), performance on adjacent pairs was above chance but comparatively low.

**Figure 2:**
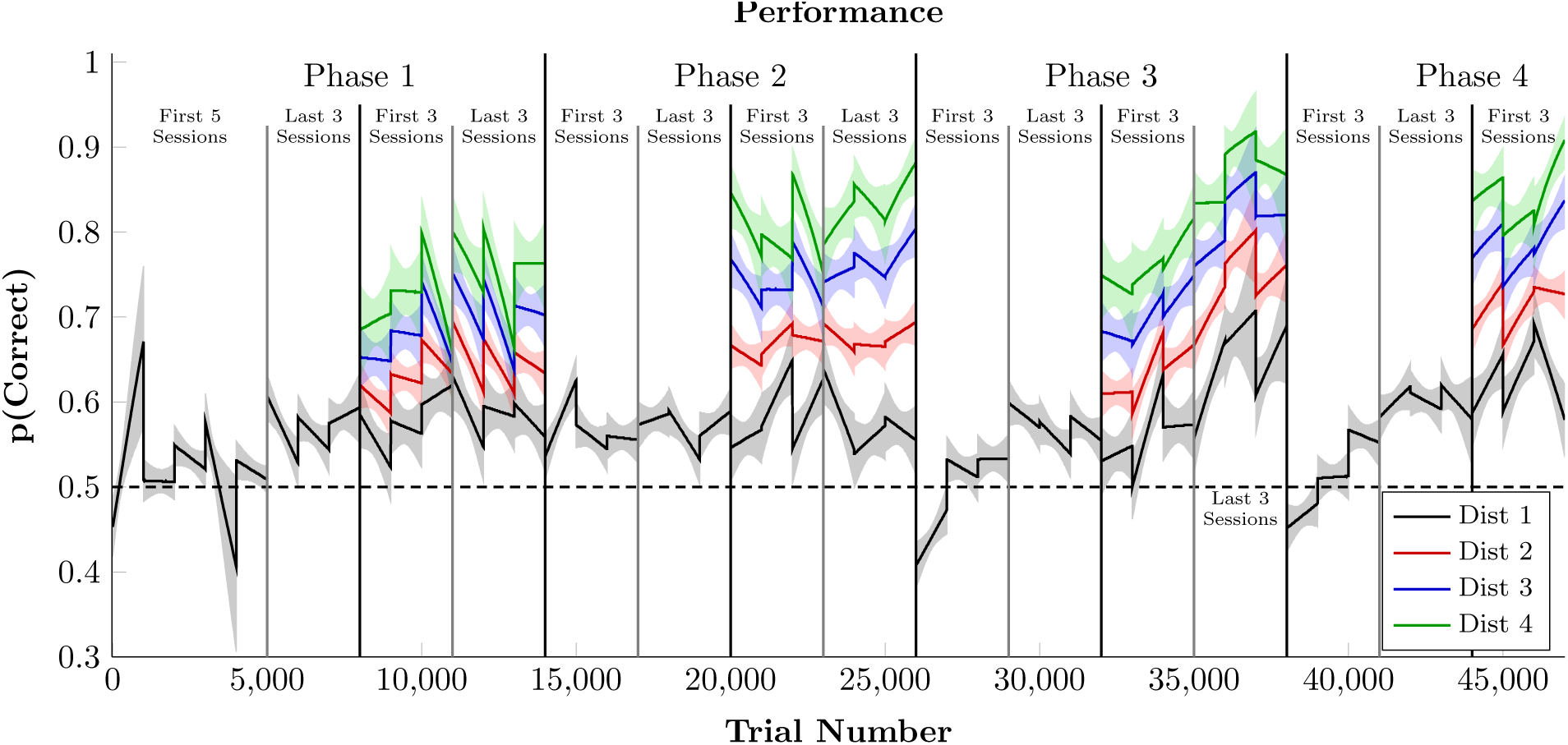
Time series analysis of task performance, divided by symbolic distance, averaged across subjects. All sessions presented adjacent pairs (black), but only all-pairs sessions included symbolic distance of 2 (red), 3 (blue), and 4 (green). Discontinuities correspond to gaps between sessions. Shaded regions represent the 99% credible interval of the estimate.

After at least 6 sessions of adjacent-only training, monkeys were tested with all category pairings. Non-adjacent pairs yield higher accuracy, even in the earliest trials of each all-pairs phase. The symbolic distance effect is clearly visible. Response accuracy was highest for the largest symbolic distance of 4 (depicted in green), whereas distance 3 (in blue) and 2 (in red) yielded intermediate response accuracies. Although exemplars changed on every trial, a distance effect appeared immediately after the transition from an adjacent-pair to an all-pair design. This suggests that subjects immediately make transitive inferences at the category level. Figure 2 incorporates data from 47 sessions for each subject. Complete data and analyses are available in the electronic supplement.

Figure 3 presents a more direct depiction of the distance effect from the logistic regressions using the mean estimated *β_D_* parameter, measured in log units. This estimates the *differential* impact of symbolic distance, independent of overall performance. A positive value for the parameter indicates the traditional distance effect, with larger values corresponding to more dramatic effects. Thus, if (hypothetically) accuracy on adjacent pairs was at chance (i.e. logistic (0.0)) and if *β_D_* = 0.2, then accuracy on a pair with a distance of two would be logistic(0.2) = 0.55, and distance three would be logistic(0.4) = 0.60, and so forth. In all but three of the sessions, the 99% credible interval of the mean (depicted by the whiskers) excludes zero. The 80% credible interval (depicted by the boxes) exclude zero for all sessions.

**Figure 3:**
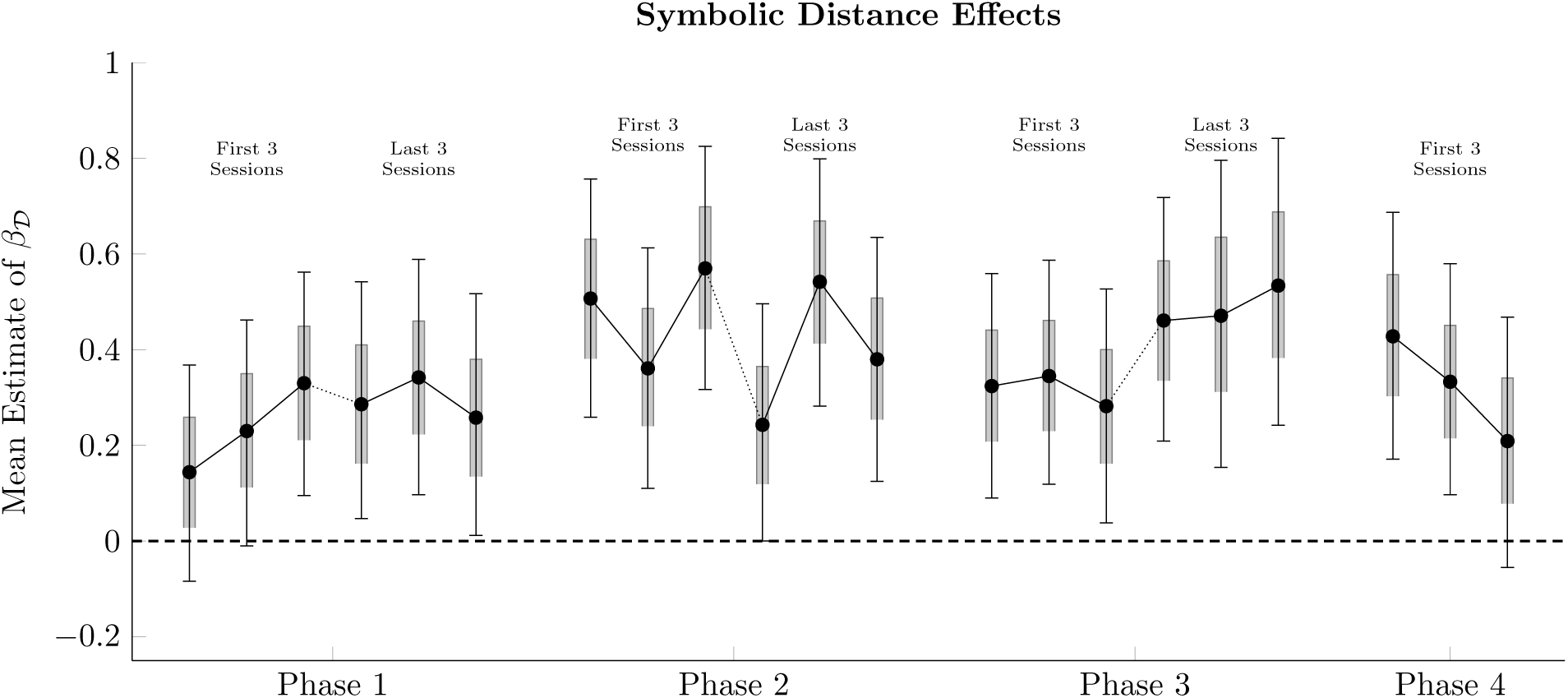
Session-by-session of the “distance effect on trial zero” parameter in the logistic regression analysis of performance (*β_D_* in equations 1 and 2) during all-pairs sessions, averaged across subjects. Since parameters are measured in log-odds units, no distance effect at transfer would correspond to a parameter value of 0.0. Whiskers represent 99% credible intervals for the estimates, while shaded intervals represent 80% credible intervals.

Reaction time was also evaluated on a per-session basis, given a log-linear model:

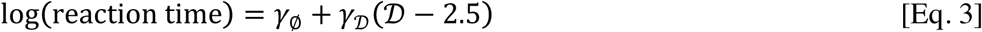

Because *γ_D_* was centered with respect to symbolic distance, the intercept *γ_ø_* can be interpreted as the mean of the log reaction times, whereas *γ_D_* is responsible for the deviation as a function of distance. This model was fit for each subject during each session.

Figure 4 plots mean parameter values across subjects for both the intercept *γ_ø_* and the distance effect *γ_D_*. Overall, reaction time increased during successive phases of the experiment, from a minimum group mean of -1.55 log seconds (0.21 s) to subsequent values reliably exceeding -0.8 log seconds (0.45 s). This suggests that subjects became more deliberative with training. Thus, despite a reliable distance effect for response accuracy, none was obtained for reaction time.

**Figure 4:**
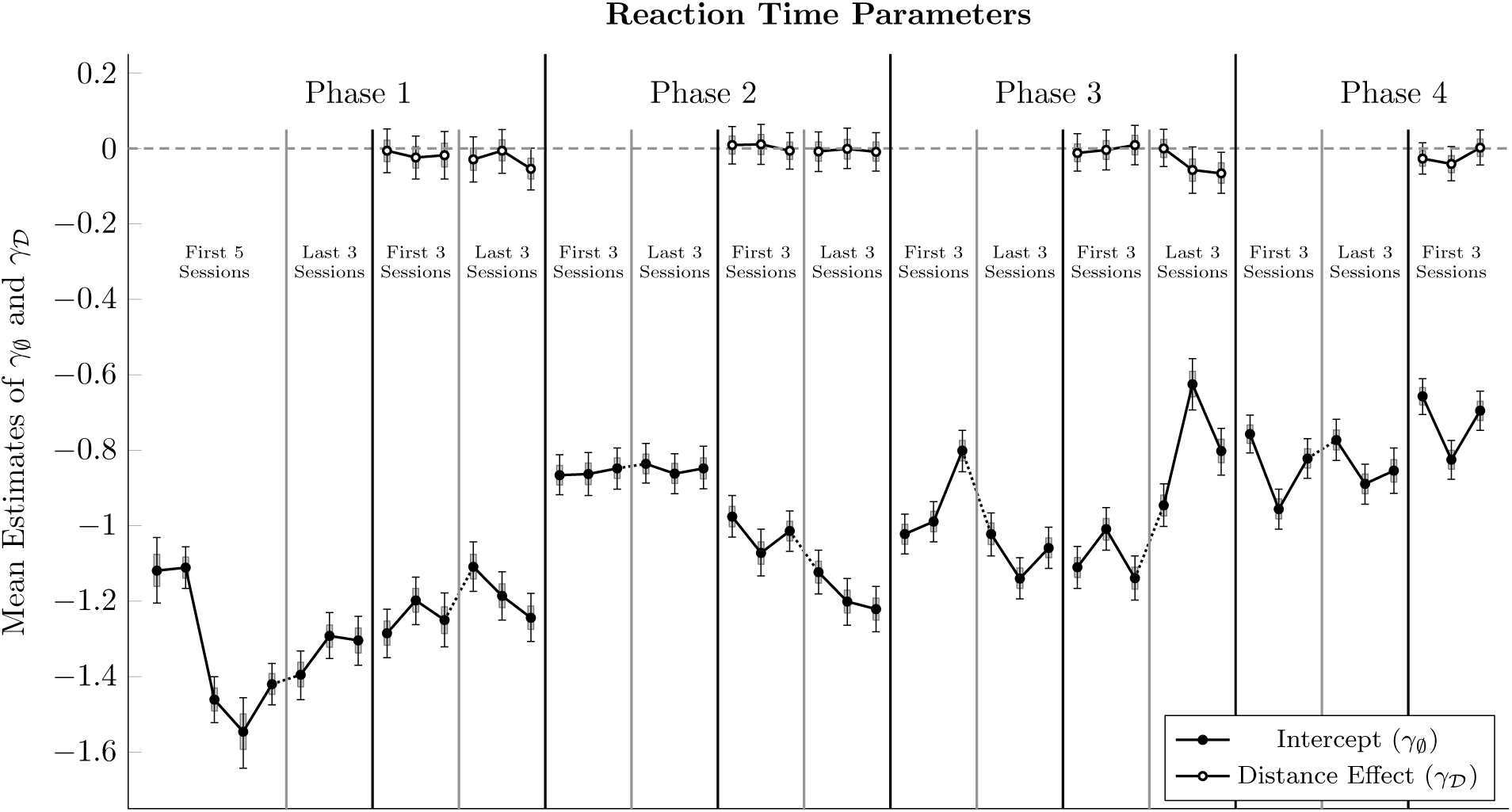
Session-by-session of the intercept parameter (*γ_ø_* in equation 3, in black) and “distance effect on trial zero” parameter (*γ_D_* in equation 3, in white) in the regression analysis of log reaction time, averaged across subjects. Values of *γ_D_* near zero indicate no differential effect on reaction time as a function of symbolic distance. Whiskers represent 99% credible intervals for the estimates, while shaded intervals represent 80% credible intervals.

## Discussion

Unlike earlier studies of category formation, we showed that rhesus macaques could be trained by a transitive inference paradigm to differentiate five perceptual categories (birds, cats, flowers, people, hoofed animals) and to learn their ordinal positions on four different implicit lists. Remarkably, the overwhelming majority of stimulus pairs were trial-unique.

Evidence of a distance effects was also obtained on the last two lists (cf. Figure 3). For both monkeys, distance 4 is shown in green, distance 3 in blue, distance 2 in red and distance 1 in black. Across all phases, response accuracy was lowest for pairs at distance 1, despite significantly more training on those pairs than pairs with symbolic distances of 2, 3, and 4. Taken together, our results show that monkeys could retain knowledge of five distinct perceptual categories, despite changes to the ordering of the categories, that they could readily update the ordering of those categories and improve their performance with experience.

Although the estimated distance effect was consistently positive (i.e. larger symbolic distances yielded higher performance), the effect size in the first two sessions of phase 1 was not statistically significant. During those sessions, subjects may still have been learning to categorize the exemplars. Alternatively, they might have had a less robust understanding of the order of the categories. However, in every subsequent transfer, subjects showed a clear and statistically significant distance effect. Past studies have shown that monkeys’ performance improves as they accrue expertise over consecutive sessions learning serial tasks (Terrace et al. 2003). Response accuracy in later stages resembled TI in other studies (e.g. Jensen et al. 2015), suggesting that given sufficient expertise, subjects were able to manipulate categories as though each was a “stimulus.”

The analysis of reaction times yielded two surprising results. Extensive training increased reaction time and, unlike response accuracy, no reliable distance effect of reaction time emerged.

These effects are likely due to the change of exemplars on every trial. While a monkey that lacks category knowledge can respond rapidly by guessing, a monkey that seeks to classify an exemplar will need more time to identify it.

Traditionally, studies of categorization in animals initially train category membership using the match-to-sample paradigm (Herrnstein 1985, Crouzet et al. 2012), a match-to-stimulus paradigm (Fabre-Thorpe 1998, Basile & Hampton 2013), or a match-to-category design (Freedman and Miller 2001). In these paradigms, subjects evaluate stimuli one at a time, a process that is highly vulnerable to a “guessing” strategy (Jensen & Altschul 2015). The categorical TI experiment is distinct from these procedures because it required subjects to evaluate two categories from ten possible pairings. Subjects not only learned to discriminate the categories, but did so while *simultaneously* learning the ordinal positions of those categories.

### Cognitive Representation Of Serial Categories

Proposals of how animals categorize stimuli can be grouped into two classes: associative learning and cognitive representation. Roberts (1996) and Lea and Ryan (1984) argued that animals ability to categorize can be explained by their reinforcement history. Because category exemplars contain particular features, they can be paired with rewards. But this interpretation raises an obvious question: What are those features? Herrnstein and Perrett (1985) questioned that interpretation in an experiment in which photographic stimuli were randomly assigned to categories without regard to their content. Pigeons were nevertheless able to learn which images belonged to which category.

A more modern cognitive approach treats a perceptual category as a “conceptual representation” (Newen & Bartels 2007). Under such a view, categorization arises from an animal’s ability to embed stimuli into a representational hierarchy, such that stimuli can both be decomposed into features and also be grouped into categories. These groupings can be defined statistically, rather than by strict rules. For example, although “humans” might consistently have two eyes, an animal would be able to categorize a human with only one eye if enough other features were consistent with those of other members of the superordinate grouping. As such, no single feature is necessary or sufficient to determine category membership. Instead, the hierarchical representation overall would permit categorization. Although such representations fall short of the abstract sophistication of language, they are nevertheless more generalized and flexible than reward associations to individual features. Categorization based on conceptual representations requires consistent, correct classification of diverse stimuli that is not based on task-related discriminative cues, and requires abstraction of stimuli that cannot be categorized by generalization of features alone.

Prior to the 1970s, TI was thought to rely on logic, thereby limiting it to humans, age seven and older, who possessed both language and the cognitive capacity to perform concrete operations (Vasconcelos, 2008). However, Bryant and Trabasso (1971) demonstrated that four-year-old children displayed TI prior to the manifestation of concrete operations, suggesting that TI depends on a more fundamental cognitive capacity. Their method of training list order by trial and error was translated to non-human animals by McGonigle and Chalmers (1977), who found evidence of TI in squirrel monkeys (*Saimiri sciureus*).

Although evidence for TI in animals is compelling, its underlying mechanism is more difficult to resolve. Some have argued that associative learning (often using some variant of the Rescorla-Wagner theory of learning) is the most parsimonious explanation for TI’s ubiquity in animals (Vasconcelos, 2008). However, there are serious problems with this argument. According to associative models, the massed presentation of a single stimulus pair (e.g. DE) should generally bias responding toward the correct item in that pair, even in other pairings where it is incorrect (e.g. BD). However, several species have demonstrated robust response accuracy despite these manipulations (Lazareva & Wasserman 2012, Jensen et al. 2016).

Both the consistent manifestation of symbolic distance effects and TI’s resistance to the effect of massed trials suggests that behavior is mediated by a cognitive representation, which is updated based upon feedback. For example, Jensen and colleagues (2015) proposed a Bayesian model in which subjects estimate probability distributions for the position of each stimulus along a spatial continuum, and judge which stimulus to select by first drawing random samples from those distributions, then selecting the stimulus whose sample yielded a lower score. Our data display symbolic distance effects and transfer effects for critical test pairs that are consistent with the predictions of a Bayesian spatial model.

That said, it would be a mistake to make too broad a claim, based on our data, about categorization and TI. All of the stimuli used in this experiment were photographic images. Although the photographs were taken at different angles, colors, and degrees of zoom, there inevitably were statistical and featural regularities among images. For example, pictures of flowers never included eyes, the absence of which could be used as a cue for that category. The present study does not rule out the possibility that subjects relied on a classifier that was tailor-made for the stimulus set, shaped by this study’s specific feedback (Jensen & Altschul, 2015). However, this does not alter our conclusions regarding serial learning. A tailor-made classifier might perform more poorly on novel stimuli than a general-purpose classifier. But in either case, the subjects would be performing TI at a level of abstraction above that of specific stimuli.

Another potential concern regarding the use of photographic stimuli is that they may be “ecologically relevant,” such that subjects might have some biological predisposition to categorize them correctly (New et al. 2007). In light of both of these concerns, a replication of our design using artificial stimuli (e.g. man-made stimuli) would be illuminating. However, we are not making any assertions about how categorization is performed, or whether it is innate or acquired. Past studies of animal categorization suggest that animals still exhibit serial learning with abstract artificial stimuli (Altschul et al. 2016), and that they are able to categorize visually degraded photographic stimuli (Basile & Hampton, 2013) and artificial stimuli (Matsukawa et al. 2004). Thus, although our own use of photographic stimuli may introduce an ecological confound, an ample literature suggests that subjects should be able to learn to categorize and serially order stimuli beyond those that are “ecologically relevant.”

## Acknowledgements

We thank Grant Spencer, Orly Morgan, David Freshwater and Carolina Montes for assistance with data collection.

## References

Altschul D, Jensen G, Terrace HS. (2016). Perceptual category learning of photographic and painterly stimuli in rhesus macaques (*Macaca mulatta*) and humans. PeerJ PrePrints, 4, e967v3.

Basile BM, Hampton RR. (2013). Monkeys show recognition without priming in a classification task. Behavioural Processes, 93, 50-61.

Bhatt RS, Wasserman EA, Reynolds Jr. WF, Knauss, KS. (1988). Conceptual behavior in pigeons: Categorization of both familiar and novel examples from four classes of natural and artificial stimuli. Journal of Experimental Psychology: Animal Behavior Processes, 14, 219-234.

Bodily KD, Katz JS, Wright AA. (2008). Matching-to-sample abstract-concept learning by pigeons. Journal of Experimental Psychology: Animal Behavior Processes, 34, 178-184.

Bryant PE, Trabasso T. (1971). Transitive inference and memory in young children. Nature, 232, 456-458.

Carpenter B, Gelman A, Hoffman M, Lee D, Goodrich B, Betancourt M, Brubaker MA, Guo J, Li P, Riddell A. (In Press). Stan: A probabilistic programming language. Journal of Statistical Software.

Chen S, Swartz KB, Terrace HS. (1997). Knowledge of the ordinal position of list items in rhesus monkeys. Psychological Science, 8, 80-86.

Crouzet SM, Joubert OR, Thorpe SJ, Fabre-Thorpe M. (2012). Animal detection precedes access to scene category. PLOS ONE, 7, e51471.

D’Amato MR, Colombo M. (1990). The symbolic distance effect in monkeys (*Cebus apella*). Animal Learning & Behavior, 18, 133-140.

Fabre-Thorpe M, Richard G, Thorpe SJ. (1998). Rapid categorization of natural images by rhesus monkeys. NeuroReport, 9, 302-308.

Freedman DJ, Riesenhuber M, Poggio T, Miller EK (2001). Categorical Representation of Visual Stimuli in the Primate Prefrontal Cortex. Science, Vol. 291, Issue 5502, 312-316.

Herrnstein RJ, Loveland DH. (1964). Complex visual concept in the pigeon. Science, 146, 549-551.

Herrnstein RJ, Perrett DI. (1985). Riddles of natural categorization [and discussion]. Philosophical Transactions of the Royal Society of London. Series B, Biological Sciences, 308, 129-144.

Jensen G. (2017). Serial Learning. In Call J (Ed.), APA Handbook of Comparative Psychology, Vol.2 (Ch. 18). Washington, DC: APA Books.

Jensen G, Altschul D, Danly E, Terrace HS. (2013). Transfer of a serial representation between two distinct tasks by rhesus macaques. PLOS ONE, 8, e70285.

Jensen G & Altschul D. (2015). Two perils of binary categorization: Why the study of concepts can’t afford true/false testing. Frontiers of Psychology, 6, Article 168.

Jensen G, Muñoz F, Alkan Y, Ferrera VP, Terrace HS. (2015). Implicit value updating explains transitive inference performance: The betasort model. PLOS Computational Biology, 11, e1004523.

Jensen G, Alkan Y, Muñoz F, Ferrera VP, Terrace HS. (2016). Transitive inference in humans and rhesus macaques after massed training of the last two list items. bioRxiv, doi:10.1101/055335.

Lazareva OF, Smirnova AA, Bagozkaja MS, Zorina ZA, Rayevsky VV, Wasserman EA. (2004). Transitive responding in hooded crows requires linearly ordered stimuli. Journal of the Experimental Analysis of Behavior, 82, 1-19.

Lazareva OF, Wasserman EA. (2006). Effect of stimulus orderability and reinforcement history on transitive inference in pigeons. Behavioural Processes, 72, 161-172.

Lazareva OF, Wasserman EA. (2012). Transitive inference in pigeons: Measuring the associative values of stimuli B and D. Behavioural Processes, 89, 244-255.

Lea SEG, Ryan ME. (1984). Feature analysis of pigeons’ acquisition of concept discrimination. In Commons ML, Herrnstein RJ, Wagner AR (Eds.), Quantitative Analyses of Behavior: Discrimination Processes (pp.239-253). Cambridge, MA: Ballinger.

Marsh HL, MacDonald SE. (2008). The use of perceptual features in categorization by orangutans (Pongo abelli).

Matsukawa A, Inoue S, Jitsumori M. (2004). Pigeon’s recognition of cartoons: effects of fragmentation, scrambling, and deletion of elements. Behavioural Processes, 65, 25-34.

McGonigle BO, Chalmers M. (1977). Are monkeys logical? Nature, 267, 694-696.

McGonigle BO, Chalmers M. (1992). Monkeys are rational! Quarterly Journal of Experimental Psychology, 45, 189-228.

New J, Cosmides L, Tooby J. (2007). Category-specific attention for animals reflects ancestral priorities, not expertise. Proceedings of the National Academy f Science USA, 104, 16598-16603.

Newen A, Bartels A (2007) Animal minds and the possession of concepts. Philosophical Psychology, 20, 283-308.

Roberts WA. (1996). Stimulus generalization and hierarchical structure in categorization by animals. In Zentall TR, Smeets PM (Eds.), Stimulus Class Formation in Humans and Animals (pp.35-54). Amsterdam: Elsevier.

Terrace HS. (1984). Simultaneous chaining: The problem it poses for traditional chaining theory. In Commons ML, Herrnstein RJ, Wagner AR (Eds.), Quantitative Analyses of Behavior: Discrimination Processes (pp.115-138). Cambridge, MA: Ballinger.

Terrace HS, Son LK, Brannon EM. (2003) Serial expertise of rhesus macaques. Psychological Science, 14, 66-73.

Terrace HS. (2005) The simultaneous chain: A new approach to serial learning. Trends in Cognitive Sciences, 9, 202-210.

Treichler FR, Raghanti MA, Van Tilburg DN. (2007). Serial list linking by macaque monkeys (*Macaca mulatta*): List property limitations. Journal of Comparative Psychology, 121, 250-259.

Vasconcelos M. (2008). Transitive inference in non-human animals: An empirical and theoretical analysis. Behavioural Processes, 78, 313-334.

Watanabe S. (2013). Preference for and discrimination of paintings by mice. PLOS ONE, 8, e65335.

